# Glucocorticoid receptor gene polymorhisms and breast cancer: significant association in cancerous tissue samples from Turkish patients

**DOI:** 10.1101/292631

**Authors:** Gürkan Akyildiz, Mustafa Akkіprіk, Ayşe Özer, Handan Kaya, Bahadіr M. Güllüoğlu, Cenk Aral

## Abstract

***Background and Aims:*** Glucocorticoid receptor (GR) has been postulated to serve an important role in normal breast and breast carcinoma. On the other hand, studies on its common polymorphisms with breast cancer are very limited. In this study, we aimed to assess the presence and frequency of 4 common polymorphisms of the GR gene in the cancerous tissue samples from breast cancer patients in the Turkish population and compare them with healthy individuals. ***Methods:*** DNA samples from 86 Turkish female breast cancer patients and 86 healthy controls analysed for BclI, Tth111I, N363S and ER22/23EK polymorphisms of GR gene by PCR-RFLP. ***Results:*** N363S polymorphism was not observed in study samples. Frequency of BclI polymorphism did not differ between cases and controls but it associated with family history of cancer. The polymorphic allele frequency of Tth111I and ER22/23EK was found to be higher in cases compared to healthy samples (p<0.05). Also, a haplotype containing polymorphic variants of these two polymorphisms found to be associated with breast cancer. ***Conclusion:*** The data presented here suggest that polymorphisms of GR gene may be an important risk factor for breast cancer predisposition.

## Introduction

Breast cancer is the leading malignancy among female population worldwide as well as in Turkey. Estimated age-standardized incidence and mortality rates of breast cancer among Turkish women are of 24.5 % and 15.7%, respectively [1]. The identification of the molecular genetic factors responsible for the development and progress of breast cancer should contribute to the prevention and treatment of the disease.

In the past decade, the role of glucocorticoids and of their receptors in the development, progress, and treatment of breast cancer have been widely investigated [2–4]. Glucocorticoids exert their effects via the glucocorticoid receptor (GR) which belongs to the nuclear receptor superfamily. Following the ligand binding, the GR translocates to the nucleus and affects gene expression of the target genes. Several studies indicate a relationship between GR expression and breast cancer [2, 3, 5–7]. For example Pan et al. [5], reported a better outcome in estrogen receptor-positive breast cancer patients with high level of GR expression and suggested a direct role of GR in determining the outcome of poor prognosis breast cancers. West et al. [6], suggested that co-expression of estrogen receptor and GR contributes to a less aggressive breast cancer. Belova et al. [8], found a significantly higher expression of GR in breast cancer patients older than 50 years old.

With regard to the polymorphisms of the GR gene (NR3C1), we found only 2 studies that are concerned with and reveal their link to breast cancer. One of them belongs to Takeo et al [9]. This article is in Japanese and we are unable to reach and read the full article. Thus, we can only assume that the authors display some data according to the abstract section as published on Pubmed. In the second one, Curran et al. investigated a polymorphic dinucleotide repeat (D5S207) situated 200 kb adjacent to the GR gene [10]. They found 6 different alleles in breast cancer patients and control samples with a significant difference. In our study, we aim at investigating the prevalence of 4 common polymorphisms of the GR gene in breast cancer patients and healthy controls. As a preliminary study, we have used DNA samples from our previous study and genotyped for BclI, Tth111I, ER22/23EK and N363S polymorphisms. The frequency of each polymorphism, and haplotypes has been compared between cases and controls and a significant difference has been found for Tth111I, ER22/23EK polymorphisms.

## Materials and methods

### Cases

In this study, we used previously isolated DNA samples from paraffin embedded cancerous tissues from Turkish sporadic breast cancer patients and from blood samples of healthy individuals. The detailed information regarding the patients has been given previously [11]. DNA samples from 86 healthy controls were also examined for the GR gene polymorphisms and compared to the patient group. The study was approved by the local research ethics committee of the School of Medicine at Marmara University.

### Genotyping

The presence of the GR polymorphisms was determined by PCR and restriction enzyme length polymorphism assay. Primers used for amplification were purchased from Integrated DNA Technologies and their sequences are given in Table I.

PCR conditions were the same for all loci of polymorphisms except annealing temperatures which are indicated in Table I. All reactions consisted of 5 min of initial denaturation at 95 °C followed by 40 cycles of 1 min denaturation at 95 °C, 1 min annealing, and 1 min elongation at 72 °C. All PCR reactions were confirmed by 2 % agarose gel electrophoresis. All PCR reagents were purchased from Fermentas.

In order to determine the ancestral or rare allele, BclI, Tth111I, MnlI, and TasI restriction enzymes were used for BclI, Tth111I, ER22/23EK, and N363S polymorphisms, respectively. All enzymes were purchased from Fermentas and reactions were done according to the manufacturer’s instructions.

Haplotype structures and their frequencies were estimated from genotype data within the linkage disequilibrium block using the Haploview version 4.2 (http://www.broadinstitute.org/haploview/haploview), which estimates haplotype by an accelerated EM algorithm [12].

### Statistics

PASW statistics 18 software was used to compare the frequencies of polymorphisms between cases and controls. The *χ*2 analysis, followed by Fisher’s exact test wherever required, was also used. Haplotype frequencies and linkage disequilibrium were analysed by Haploview software as noted above. A difference was determined to be statistically significant when p<0.05.

## Results and Discussion

DNA samples from 86 Turkish female breast cancer patients were included in this study retrospectively. Demographic data and clinical/pathological findings of the patients were recorded from their files, but some clinical data were unavailable for some patients. The mean age of patients was 60 ± 12. Among them, 20 patients were younger than or around 50 years old and the rest was older. With regard to the receptor expression status, 41 and 34 patients were positive for estrogen and progesterone receptor, respectively. The ER status of 31 patients and the PR status of 33 patients were not available. Some 58 patients were recorded as postmenopausal at the time of diagnosis. The menopausal status of 7 patients was not available in patients records. The number of patients with and without family history for breast and/or ovary cancer was of 10 and of 64, respectively. Tumor size, presence of axillary invasion, metastatic nodule number, and stage of cases were also recorded and compared between genotypes as given below when available.

### BclI (rs41423247)

BclI polymorphism is located in intron 2, close to 3ˊ of exon 2 (Fig. 1, A). Since it does not involve a regulatory or splicing region, it has been defined to be in linkage disequilibrium with other polymorphisms. DNA samples from 73 patients and 86 controls were successfully amplified and analysed for *BclI* polymorphism. Frequencies of each genotype were given in Table 2. As it can be seen in this table, frequencies of genotypes were similar in breast cancer samples when compared to those of healthy individuals (*p*=0.331). Clinical parameters of patients with their genotypes are given in Table 3. None of the clinical parameters is associated with this polymorphism except family history. The percentage of heterozygosity is relatively higher in family history positive patients (*p*=0.022). Only 10 cases had positive family history and it is not possible to generalise this finding without further studies, including a larger cohort of cases. A related study with other breast cancer susceptibility genes might provide a proper explanation for our findings. According to 1000 genomes project, minor allele frequency is 0.20, 0.26, 0.24, 0.38, and 0.21 in African, American, East Asian, European and South Asian populations, respectively [13]. It is 0.297 in our healthy group and slightly lower than European data. On the other hand, polymorphic allele frequency is close to our previous data from Turkish individuals [14].

**Table 1.**
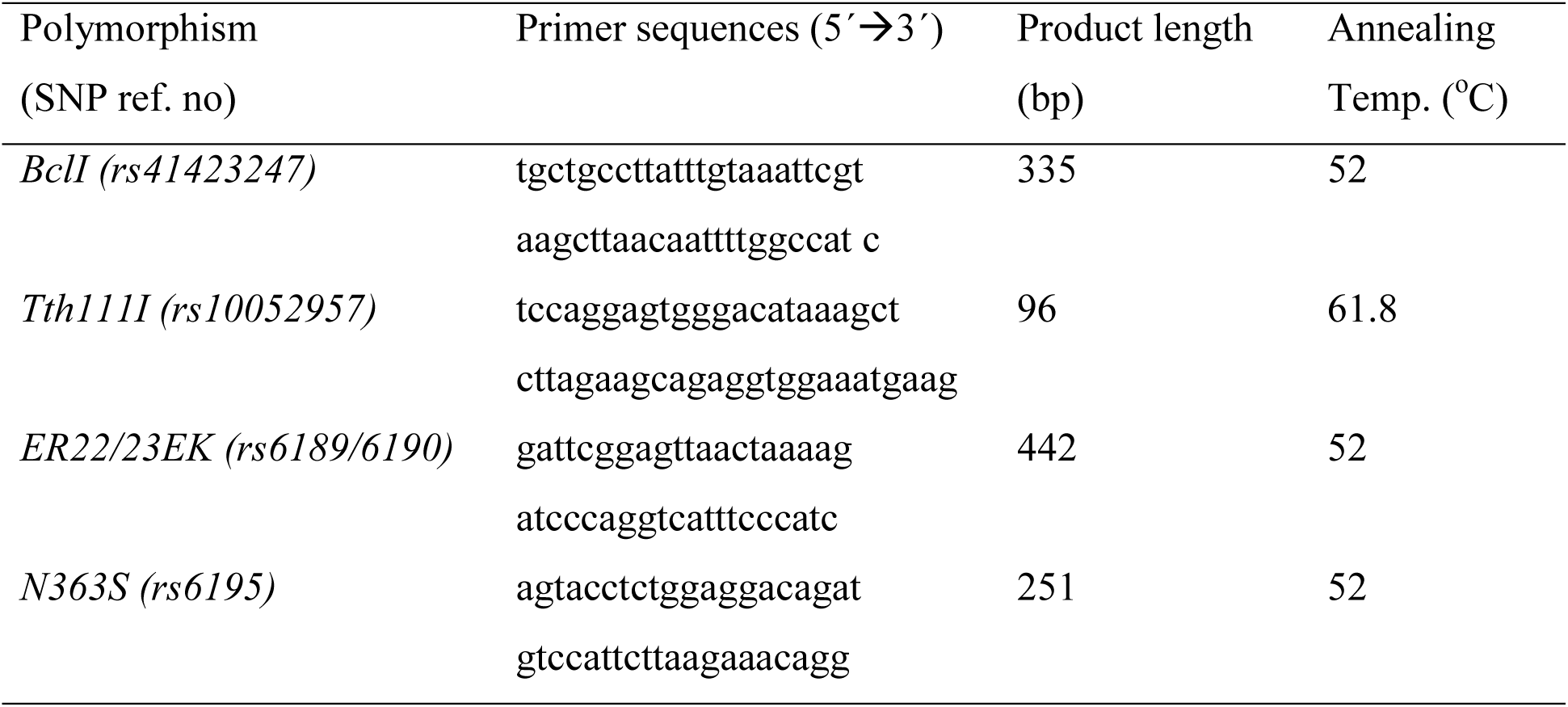
Primer sequences, product lengths and annealing temperatures used for amplification

**Table 2.** The genotype and allele frequencies of GR gene polymorphisms (* denotes ancestral allele).

|  |  | Cases, n (%) | Controls, n (%) | *p* value |
| --- | --- | --- | --- | --- |
| BclI polymorphism |  |  |  |  |
| Genotype | CC | 37 (50.7) | 40 (46.5) | 0.331 |
|  | CG | 35 (47.9) | 41 (47.7) |  |
|  | GG | 1 (1.4) | 5 (5.8) |  |
| Allele | C* | 109 (74.7) | 121 (70.3) | 0.392 |
|  | G | 37 (25.3) | 51 (29.7) |  |
| Tth111I polymorphism |  |  |  |  |
| Genotype | CC | 13 (15.7) | 31 (36.0) | 0.009 |
|  | CT | 54 (65.1) | 45 (52.3) |  |
|  | TT | 16 (19.2) | 10 (11.7) |  |
| Allele | C* | 80 (48.2) | 107 (62.2) | 0.010 |
|  | T | 86 (51.8) | 65 (37.8) |  |
| ER22/23EK polymorphism |  |  |  |  |
| Genotype | GG | 40 (85.1) | 85 (98.8) | 0.001 |
|  | GA | 7 (14.9) | 1 (1.2) |  |
|  | AA | 0 (0) | 0 (0) |  |
| Allele | G* | 87 (92.6) | 171 (99.4) | 0.002 |
|  | A | 7 (7.4) | 1 (0.6) |  |

**Table 3.** Correlations with GR gene polymorphisms and patient’s clinical parameters

| Clinical parameters |  | BclII polymorphism |  |  |  | Tth111I polymorphism |  |  |  | ER22/23EK polymorphism |  |  |  |
| --- | --- | --- | --- | --- | --- | --- | --- | --- | --- | --- | --- | --- | --- |
|  |  | CC | CG | GG | *p* | CC | CT | TT | *p* | GG | GA | AA | *p* |
| Age | ≤ 50 | 12 | 7 | 0 | .406 | 5 | 11 | 4 | .390 | 9 | 3 | 0 | .254 |
|  | > 50 | 25 | 28 | 1 |  | 8 | 43 | 12 |  | 31 | 4 | 0 |  |
| Grade | I | 7 | 7 | 0 | .748 | 5 | 8 | 6 | .234 | 7 | 1 | 0 | .725 |
|  | II | 20 | 17 | 1 |  | 5 | 28 | 9 |  | 22 | 3 | 0 |  |
|  | III | 10 | 7 | 0 |  | 3 | 14 | 1 |  | 9 | 3 | 0 |  |
| ER status | + | 18 | 19 | 0 | .747 | 7 | 27 | 6 | .536 | 20 | 2 | 0 | .322 |
|  | - | 7 | 6 | 0 |  | 3 | 6 | 3 |  | 7 | 2 | 0 |  |
| PR status | + | 14 | 16 | 0 | .679 | 7 | 21 | 5 | .490 | 16 | 2 | 0 | .592 |
|  | - | 9 | 8 | 0 |  | 3 | 9 | 5 |  | 9 | 2 | 0 |  |
| Menopausal status | Pre | 11 | 8 | 0 | .790 | 5 | 12 | 4 | .496 | 11 | 3 | 0 | .465 |
|  | Post | 26 | 22 | 1 |  | 7 | 36 | 12 |  | 27 | 4 | 0 |  |
| Tumor size | T1 | 15 | 12 | 0 | .381 | 3 | 19 | 7 | .056 | 17 | 2 | 0 | .673 |
|  | T2 | 11 | 16 | 1 |  | 6 | 19 | 5 |  | 14 | 4 | 0 |  |
|  | T3 | 6 | 2 | 0 |  | 4 | 2 | 1 |  | 3 | 1 | 0 |  |
|  | T4 | 2 | 0 | 0 |  | 0 | 1 | 2 |  | 2 | 0 | 0 |  |
| Nodule number | n0 | 14 | 13 | 0 | .348 | 5 | 18 | 6 | .967 | 15 | 3 | 0 | .872 |
|  | n1 | 8 | 10 | 0 |  | 3 | 13 | 4 |  | 10 | 1 | 0 |  |
|  | n2 | 5 | 5 | 0 |  | 3 | 6 | 2 |  | 4 | 1 | 0 |  |
|  | nx | 7 | 3 | 1 |  | 2 | 5 | 3 |  | 7 | 2 | 0 |  |
| Lymphatic invasion | yes | 13 | 15 | 0 | .392 | 6 | 19 | 6 | .713 | 14 | 2 | 0 | .626 |
|  | no | 14 | 10 | 0 |  | 3 | 17 | 6 |  | 13 | 3 | 0 |  |
| Family history | yes | 2 | 8 | 0 | .022 | 1 | 6 | 3 | .725 | 3 | 2 | 0 | .126 |
|  | no | 32 | 22 | 0 |  | 11 | 39 | 13 |  | 33 | 5 | 0 |  |
| Stage | 1 | 9 | 9 | 0 | .937 | 2 | 13 | 4 | .558 | 12 | 2 | 0 | .649 |
|  | 2 | 12 | 13 | 0 |  | 4 | 18 | 6 |  | 12 | 3 | 0 |  |
|  | 3 | 6 | 5 | 0 |  | 4 | 7 | 1 |  | 5 | 1 | 0 |  |

**Figure 1.**
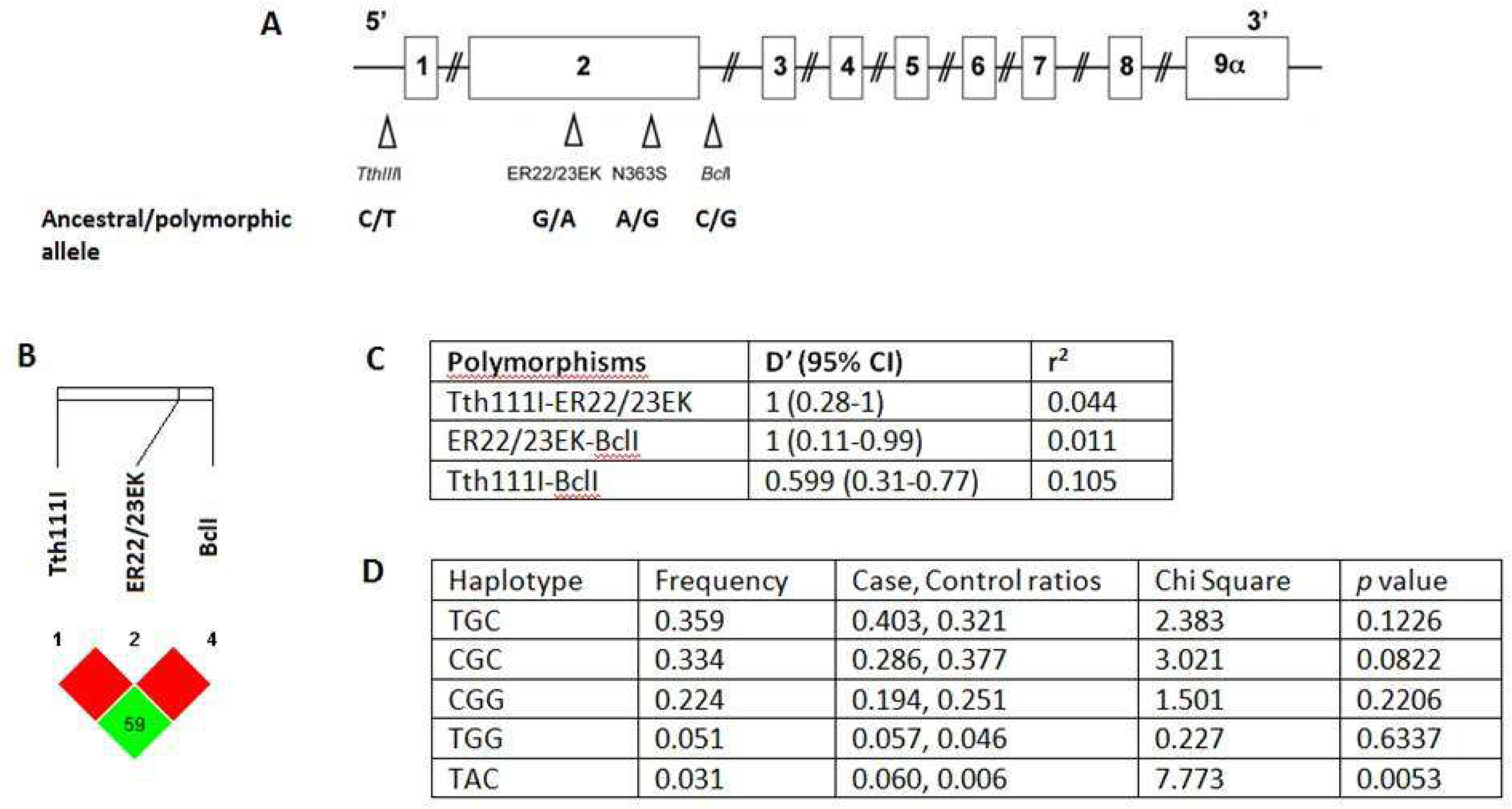
The polymorphism and haplotype analysis. A, Schematic description of glucocorticoid receptor gene and positions of polymorphisms analysed (modified from ref. [14]). B, LD plot of the investigated polymorphisms. C, a pairwise measure of linkage disequilibrium of polymorphisms with D’statistics and correlation coefficient (r^2^) values. D, haplotype patterns, their frequencies, and association with breast cancer risk.

### Tth111I (rs10052957)

Tth111I polymorphism is located 6305 bp upstream of the initiation codon and is considered to be related to promoter activity [15]. Samples from 83 breast cancer patients and 86 healthy individuals were analysed for Tth111I polymorphism. Allele frequencies were calculated from genotype data as it is given in Table 2, and it is revealed that the polymorphic allele frequency is significantly higher in patients than in controls (51.8% and 37.8%, respectively) (*p*=0.010). Polymorphic allele frequency of our healthy group (0.378) is similar to previous data from Turkish population [14], and close to European population data [13]. There were no significant correlations found between genotype frequencies and clinical variables of the patients (Table 3). Tth111I polymorphism was reported to be linked partially to ER22/23EK polymorphism [16]. In parallel, all ER22/23EK carriers of our study group also carry at least one rare allele of Tth111I polymorphism. But not all Tth111I carriers carry the ER22/23EK rare variant.

### ER22/23EK (rs6189/6190)

The polymorphism is located in the transactivation domain of the receptor gene. This polymorphism has been associated with relative glucocorticoid resistance and with healthier metabolic profile [17]. ER22/23EK polymorphism was assessed in a total of 47 patients and 86 healthy individuals. None of the samples had homozygous polymorphic genotype. This polymorphism is not found in African and South Asian populations and very low in European populations (3%) [13]. Heterozygous GA genotype was found in 7 (14.9%) patients and 1 (1.2 %) healthy individual with significant difference (*p*=0.001). All rare variant carriers also carry Ttth111I variant as noted above. As in other polymorphisms, there were no associations with clinical parameters (Table 3).

### N363S (rs6195)

This polymorphism is located in codon 363 of exon 2. It yields an amino acid alteration from asparagine (N) to serine (S) and is associated with increased sensitivity to glucocorticoids. All of the analysed samples from 63 patients and 86 healthy individuals consisted of ancestral AA genotype. Polymorphic G allele was not found. This is consistent with population data from other continents except European (%2) [13].

### Haplotype analysis

In order to determine the possible haplotypes and their association with breast cancer risk, we determined linkage disequilibrium between each polymorphism. In this analysis, N363S was excluded due to the single allele determined in the whole study population. The data are given in Figure 1. As it is seen in this figure, the neighbouring polymorphisms show stronger linkage disequilibrium with higher D’ values; however, the lower boundary CI values are below 0.7 as suggested by Gabriel et.al. We determined 5 haplotypes and also their association with breast cancer risk. Only a single haplotype showed statistically significant difference between cases and controls (*p*=0.0053). This haplotype consists of polymorphic alleles of Tth111I and ER22/23EK polymorphisms (i.e. T and A) and the ancestral allele of BclI polymorphism (i.e. C). As ER22/23EK and Tth111I polymorphisms are revealed to be linked, this result also shows the combined effect of these 2 polymorphisms.

## Conclusion

There are many risk factors involved in the development of breast cancer, including the family history of cancer, BRCA gene mutations, reproductive history, obesity, alcohol, stress, etc. Recent studies show that glucocorticoids are involved in the physiology and pathology of the breast [2]. Also, the the glucocorticoid synthesis in response to stress and its role in metabolic profiling make it an important contributor to breast cancer development and this contribution is complex. In this study, we have disclosed the direct link that exists between the common GR polymorphisms and breast cancer. There is a very limited number of studies investigating the relationship between GR polymorphisms and breast cancer. This increases the importance of the data that we have obtained in our study. ER22/23EK and Tth111I polymorphisms are significantly different between patient group and control group. As a retrospective study, we were not able to contact patients to obtain blood samples and we had to perform all analyses at cancerous tissues. This may be the most important limitation of the current study. We suggest this contribution to be further verified and clarified in the concept of some larger cohort studies and it may be relevant to validate these polymorphisms in blood samples of breast cancer patients if it is cancer tissue-specific or germline. In any case, ie. cancer tissue spesific or germline effect of the GR polymorphism, it will be an important contribution to the current knowledge of breast cancer literature.

## Acknowledgements

Authors thank D.Y.Sirin (Namik Kemal University, Department of Molecular Biology and Genetics) and P. Golban (English Language and Literature Division of Namik Kemal University) for their careful revision of the manuscript. This study is partly supported by Namik Kemal University, Scientific Research Projects Commission (NKUBAP.00.10.YL.10.28).

